# Laboratory yeast strains are highly diverse in lipid metabolism

**DOI:** 10.1101/2025.02.13.638058

**Authors:** Mike F. Renne, Richard Bachmann, Christian Klose, Thomas Hentrich, Julia M. Schulze-Hentrich, Robert Ernst

## Abstract

The impact of the genetic background on the lipidome of yeast strains remains underexplored. This study systematically compares the lipidomes of five commonly used laboratory yeast strains: BY4741, W303, D273-10B, RM11-1a, and CEN.PK2-1c. Shotgun lipidomics reveals significant variations in lipid class and acyl chain composition down to the level of molecular species. Notably, the most abundant lipid class differed between the strains: phosphatidylinositol (PI) lipids are predominant in BY4741, while phosphatidylethanolamine (PE) lipids are in D273. Ergosterol esters, which are the storage form of the major yeast sterol ergosterol, are at higher levels in all strains other than BY4741, correlating with a low gene expression of lipid metabolic enzymes Hmg1 and Are2 in BY4741. Despite these lipidomic differences, transcriptomic analysis did not show significant changes in most genes related to lipid metabolism, suggesting post-transcriptional modifications, protein abundance, and metabolic flux as potential regulatory mechanisms. This study underscores the complexity of lipidome regulation and the need for further investigation into the underlying mechanisms.

## Introduction

Lipids are essential molecules with a primary role as bulk constituents of biological membranes. In addition, lipids store metabolic energy and contribute to signal transduction. Lipids have been acknowledged as important dynamically regulated molecules with a fundamental role in cell biology, yet compared to other biomolecules such as proteins and nucleotides, their individual and collective cellular functions are less well understood [1–3]. The cellular lipid composition (*i.e.* the lipidome) is highly complex, and lipid analytical studies in various model organisms have identified tens of thousands of different structurally distinct lipids [3,4].

The budding yeast *S. cerevisiae* has been - and continues to be - an important model organism in many fields, including genetics, cell biology, and metabolism [5]. The early publication of the yeast genome [6], the ease of genetic manipulation and systematic mutant libraries [7,8], well curated databases with genetic information [9], and many other factors have contributed to the fact that yeast is arguably the best understood model eukaryote [5]. Budding yeast has been an attractive model organism to study the regulation of lipid metabolism, as its relatively simple yeast acyl chain composition makes the lipidome far less complex than that of mammalian cells, greatly simplifying lipid analysis. Yeast research has identified and dissected numerous lipid metabolic pathways, which are highly conserved amongst eukaryotes [10–12], allowing the translation of observations from yeast to higher eukaryotes.

Using mass spectrometry-based lipidomics, the lipid composition of *S. cerevisiae* has been described in great detail [13–15]. Notably, the lipidome of budding yeast is highly variable in response to growth phase and culture conditions such as the temperature, the choice of rich or defined media, and the carbon source [13–15]. Furthermore, the lipid composition is impacted by cellular and organelle stress induced by proteotoxic agents [16,17], or starvation of phosphate or carbon sources [18]. For most lipid metabolic enzymes, the regulation is elusive. How precisely cells rewire their metabolism to create these highly differing lipidomes remains unknown. Comparative lipidomic studies across different yeast species have provided fundamental insights into lipid metabolism, adaptation, and evolution [19]. However, while such studies highlight the broader diversity of lipid composition across evolutionary lineages, the extent of lipid heterogeneity within a single yeast species remains unexplored.

Currently, the field uses a plethora of laboratory yeast strains that have been obtained from natural isolates or via crossing of various background strains [20]. Large differences between wild type yeasts and genetic backgrounds have been observed, despite all strains being of the *S. cerevisiae* species. Genome-wide studies between natural isolates and laboratory strains have revealed that strains differ from each other by thousands of single nucleotide polymorphisms (SNPs), structural variations and even polyploidy [21–23]. The number of SNPs between laboratory strains and the reference genome (S288C) can range to thousands (in example ∼3.2 SNPs/kb for Σ1278b), which is comparable to the genetic variation observed between individual humans [24]. Many labs and/or topical fields have selected one or several yeast background strains as their standard model or favourite strain to study. For example, the BY4741 strain is widely used in functional genomics due to its well-characterized deletion library [20,25], while W303 has become a common choice in mitochondrial research, despite no systematic comparison of mitochondrial function between across strains. Likely due to the genetic variation between strains, different yeast genetic backgrounds show phenotypic variation to a large extent [22,26,27], ranging from sensitivity to chemicals all the way to synthetic lethal genetic interactions [28] and the essentiality of genes [24,29,30]. Notably, several studies have reported strain background dependent variations of phenotypes related to lipid metabolism [31–34] and have indicated differences in the lipid composition between strains [34,35]. However, how wild type strain backgrounds affect the lipidome has not been systematically addressed.

Here, we compare the transcriptome and lipidome of five selected wild-type laboratory strains of budding yeast. Our lipidomics analysis reveals striking differences in lipid class composition, total acyl chain composition, and lipid species composition between strains. Notably, the most abundant lipid class varies among laboratory strains and depending on the strain background the lipid class can be phosphatidylinositol (PI), phosphatidylcholine (PC), or phosphatidylethanolamine (PE). Transcriptome profiling identified substantial differences in gene expression between strains and we identified strain-specific GO-term enrichments consistent with their common laboratory use. We found notable variation in genes involved in the biosynthesis of ergosterol and ergosterol esters, consistent with strain-specific differences in ergosterol ester content. Surprisingly, while some transcriptomic changes aligned with lipidomic variations, the majority of lipidomic differences could not be explained by transcriptional variation. Our findings underscore the importance of the strain background in shaping the lipidome and unlocks the potential for new insights from repeating previous genetic screens in different strains.

## Results

### Selection of S. cerevisiae laboratory strains

To identify differences in the lipidome of different strains of budding yeast, we set out to compare the standard laboratory strain BY4741 (BY in short) to other, widely used ones. BY4741 is derived from S288C [36], which was the first eukaryote to be completely genomically sequenced yielding the reference yeast genome. BY4741 (MATa haploid), BY4742 (MAT*⍺* haploid) and BY4743 (diploid) were used as the background strains for the yeast deletion library (EuroFAN II project), and have been the strains of choice for large scale genetic studies and genome-wide screens [25,37–40]. In addition, epitope-tagged strain collections (GFP, TAP-tag, SWAT etc.) have been established in the BY background that have been used in microscopy screens [41] and mass spectrometry based studies [42–44].

For selecting the appropriate strains to compare BY with, we used data in the Saccharomyces Genome Database (SGD) Variant Viewer [45], which contains sequence similarity scores for the open reading frames of 12 widely used *S. cerevisiae* strains [46]. S288C was used in the systematic genome sequencing project that yielded the yeast reference genome, thus in literature most genetic comparisons are made to the S288C reference. BY4741 is the MATa haploid of the BY4743 diploid that was used in the systematic deletion project, which is derived from S288c [36]. Thus, BY4741 is isogenic to S288C [46]. As BY4741 is used as the reference strain in our study S288C was excluded. From the remaining 11 strains, we included 4 strain backgrounds across the board of genetic variation: W303, D273-10B, RM11-1a, and CEN.PK [45] (**Figure 1A**). (1) W303 was derived from S288C by Rodney Rothstein in a series of strain crosses but the exact ancestry of W303 is not known. It was selected to retain the prime desirable characteristics of S288C, but in addition to sporulate well and to transform with high efficiency. It shows the highest similarity, sharing > 85% of the genome with S288C [45] yet it had 7 ORFs less than S288C. It is widely used in mitochondrial studies, and the W303-derived strain W303-K6001 is used for ageing research. (2) D273-10B (D273 in short) originates from the lab of Fred Sherman and was selected for high respiration rate and resistance to glucose repression to be utilized in mitochondrial studies. Consistently, it has a low frequency of mitochondrial DNA loss (rho-penotype). (3) RM11-1a (RM11 in short) is a haploid derivative from a natural isolate from a California vineyard collected by Robert Mortimer. The sequence divergence between RM11 and S288C is between 0.5% and 1.0%. (4) The CEN.PK strain (CEN in short) was generated by Michael Ciriacy and Karl-Dieter Entian in a German consortium aiming to study metabolic fluxes in yeast [47]. CEN.PK strains are widely used in systems biology studies, and showed the lowest similarity to S288C in the SGD Variant Viewer. Between CEN.PK and the S288C reference, for example, there are > 20.000 SNPs, of which two thirds are in ORFs altering the sequence of > 1400 proteins [45]. Moreover, CEN.PK and BY4741 differ most in the number of ORFs (5379 and 5404, respectively).

**Figure 1.**
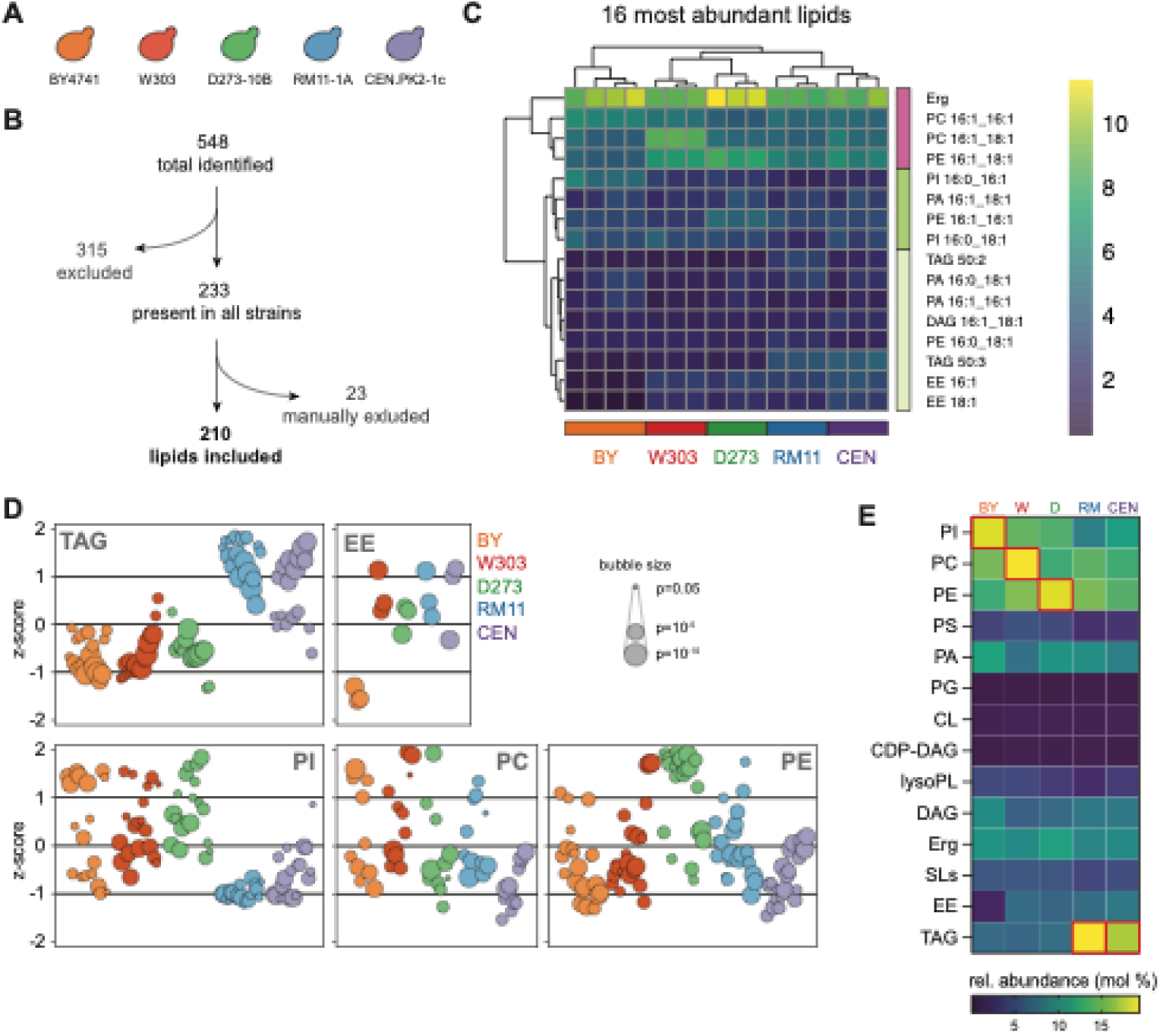
Variation in the lipid composition between different wild type yeast strains. (A) Overview of strains included in the study. (B) Schematic overview of the inclusion-criteria of detected lipids for comparison between strains. (C) Heatmap and hierarchical clustering of the 16 most abundant lipid species identified in all samples. The 16 most abundant lipids together comprise over 50% of the lipid abundance. Underlying data can be found in Table S2. (D) Bubble plot of lipid z-scores for all 5 strains tested for indicated lipids. Each bubble represents a single lipid species, and bubbles are colour coded per lipid class as indicated. Size of the bubbles corresponds to the adjusted p-value (ANOVA with correction for multiple testing). Only lipid species with an adjusted p-value < 0.05 are shown. (E) Heatmap of lipid class composition. Red squares indicate the most abundant lipid for each strain. All lysoglycerophospholipids (lysoPL) are grouped together.

### Lipidomics analysis reveals large lipidome heterogeneity between strains

To comprehensively study the lipidome of the selected strains, we employed high resolution mass spectrometry-based shotgun lipidomics [13,14]. Cells were cultured in synthetic defined glucose media and cultured to early exponential growth to warrant optimal cell growth and to prevent nutrients in the media being limiting. A total of 548 individual lipid species were detected in the total dataset (**Table S1**). As we aim to compare the analysed strains, we chose to include only lipid species detected in all samples leaving a total of 233 species. As *S. cerevisiae* naturally only synthesizes fatty acids with an even number of carbon atoms and one or no unsaturation [48], we manually excluded 23 phospholipids that contain odd chain- or polyunsaturated acyl chains and had extreme low abundance (<0.01 mol% of total), leaving a final dataset of 210 lipid species (**Figure 1B, Table S2**). Although these inclusion criteria facilitate a fair comparison of the lipidome between the strains, it does exclude lipids that may be present or absent in some strains and could provide interesting indications. For example, PC 16:0/16:0 is a low abundant but detectable lipid in BY [13,49] and was found in all strains except for RM (**Table S1**). Moreover, EE 14:0 could not be detected in BY (**Table S1**) despite previous reports [15,16], but it was present in the 4 other strains (W303, D273, RM11 and CEN).

For identifying the main differences between the strains, we conducted an analysis of the total dataset consisting of the 217 identified lipid species (**Figure S1A**). A striking variation of the lipid composition was observed between the strains, which was particularly visible for the 16 most abundant lipids that together make approximately 50% of the total lipid abundance (**Figure 1C, S1B**). Ergosterol (Erg) is the major sterol in yeast and the most abundant lipid species [13]. Although Erg levels in the ER are known to be tightly maintained, the strains show variation in the total Erg content ranging from 8.4 mol% in RM11 to 10.9 mol% in D273. Striking differences in abundance were also observed for the major glycerophospholipid species in yeast, particularly for BY, D273 and RM. BY has higher levels of the major PI species PI 16:0_16:1 and PI 16:0_18:1 compared to the other 4 strains, whereas W303 is more abundant in PC 16:1_18:1 and D273 has higher abundance of phosphatidylethanolamine (PE) 16:1_18:1 and PE 16:1_16:1 species. RM11 and CEN stand out particularly by higher abundance of TAG 50:3 than the other strains. Finally, BY has strikingly lower levels of the major ergosterol esters (EE) EE 16:1 and EE 18:1 compared to the other strains, leading to a reduction in the total EE abundance in BY4741 (forthcoming **Table 1**).

**Table 1.**
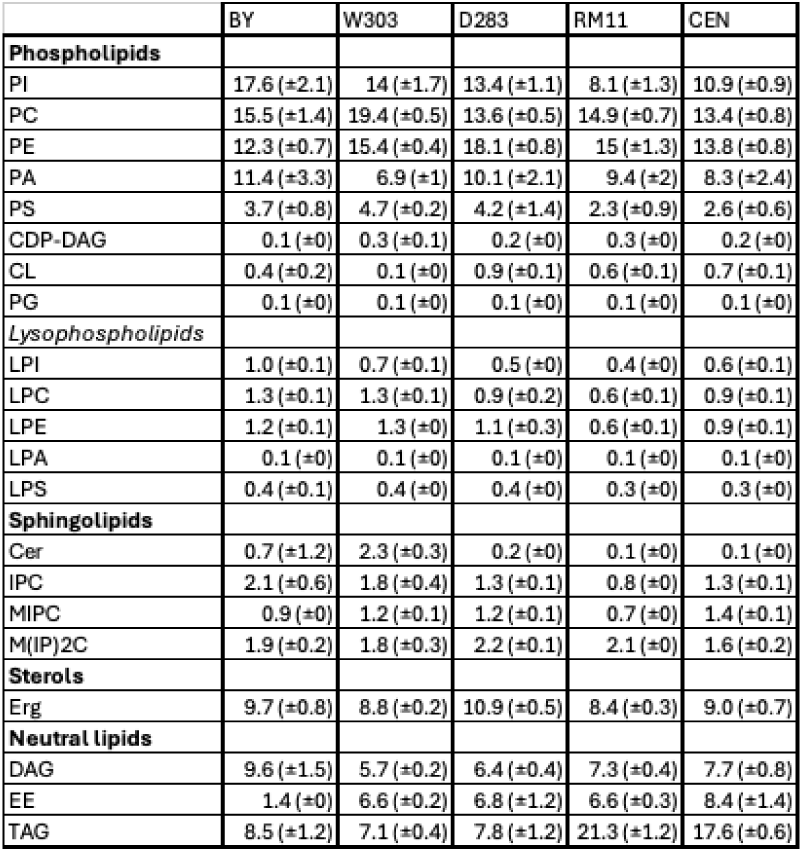
Lipid class composition of the analyzed strains. Data is given as the relative abundance (mol% as mean ± SD).

|  | BY | W303 | D283 | RM11 | CEN |
| --- | --- | --- | --- | --- | --- |
| **Phospholipids** |  |  |  |  |  |
| PI | 17.6 (±2.1) | 14 (±1.7) | 13.4 (±1.1) | 8.1 (±1.3) | 10.9 (±0.9) |
| PC | 15.5 (±1.4) | 19.4 (±0.5) | 13.6 (±0.5) | 14.9 (±0.7) | 13.4 (±0.8) |
| PE | 12.3 (±0.7) | 15.4 (±0.4) | 18.1 (±0.8) | 15 (±1.3) | 13.8 (±0.8) |
| PA | 11.4 (±3.3) | 6.9 (±1) | 10.1 (±2.1) | 9.4 (±2) | 8.3 (±2.4) |
| PS | 3.7 (±0.8) | 4.7 (±0.2) | 4.2 (±1.4) | 2.3 (±0.9) | 2.6 (±0.6) |
| CDP-DAG | 0.1 (±0) | 0.3 (±0.1) | 0.2 (±0) | 0.3 (±0) | 0.2 (±0) |
| CL | 0.4 (±0.2) | 0.1 (±0) | 0.9 (±0.1) | 0.6 (±0.1) | 0.7 (±0.1) |
| PG | 0.1 (±0) | 0.1 (±0) | 0.1 (±0) | 0.1 (±0) | 0.1 (±0) |
| **Lysophospholipids** |  |  |  |  |  |
| LPI | 1.0 (±0.1) | 0.7 (±0.1) | 0.5 (±0) | 0.4 (±0) | 0.6 (±0.1) |
| LPC | 1.3 (±0.1) | 1.3 (±0.1) | 0.9 (±0.2) | 0.6 (±0.1) | 0.9 (±0.1) |
| LPE | 1.2 (±0.1) | 1.3 (±0) | 1.1 (±0.3) | 0.6 (±0.1) | 0.9 (±0.1) |
| LPA | 0.1 (±0) | 0.1 (±0) | 0.1 (±0) | 0.1 (±0) | 0.1 (±0) |
| LPS | 0.4 (±0.1) | 0.4 (±0) | 0.4 (±0) | 0.3 (±0) | 0.3 (±0) |
| **Sphingolipids** |  |  |  |  |  |
| Cer | 0.7 (±1.2) | 2.3 (±0.3) | 0.2 (±0) | 0.1 (±0) | 0.1 (±0) |
| IPC | 2.1 (±0.6) | 1.8 (±0.4) | 1.3 (±0.1) | 0.8 (±0) | 1.3 (±0.1) |
| MIPC | 0.9 (±0) | 1.2 (±0.1) | 1.2 (±0.1) | 0.7 (±0) | 1.4 (±0.1) |
| M(IP)2C | 1.9 (±0.2) | 1.8 (±0.3) | 2.2 (±0.1) | 2.1 (±0) | 1.6 (±0.2) |
| **Sterols** |  |  |  |  |  |
| Erg | 9.7 (±0.8) | 8.8 (±0.2) | 10.9 (±0.5) | 8.4 (±0.3) | 9.0 (±0.7) |
| **Neutral lipids** |  |  |  |  |  |
| DAG | 9.6 (±1.5) | 5.7 (±0.2) | 6.4 (±0.4) | 7.3 (±0.4) | 7.7 (±0.8) |
| EE | 1.4 (±0) | 6.6 (±0.2) | 6.8 (±1.2) | 6.6 (±0.3) | 8.4 (±1.4) |
| TAG | 8.5 (±1.2) | 7.1 (±0.4) | 7.8 (±1.2) | 21.3 (±1.2) | 17.6 (±0.6) |

Clustering of strains based on the lipidome (**Figure 1C, S1A**) combined with sample correlation analysis (**Figure S1C**) and principal component analysis (PCA, **Figure S1D**) revealed 4 major clusters, in which BY4741, W303 and D273 cluster individually, while RM11 and CEN.PK cluster together. Notably, while BY4741 is arguably the most widely used model strain in lipid metabolism research, this strain is most distinct from the other four strains in clustering, PCA and correlation analysis (**Figure 1C, S1C,D**). Hierarchical clustering of the lipids in the heatmap revealed two major clusters in the lipidomics dataset: a small cluster with only 4 lipids and a large cluster with all other lipids, which is further divided into sub-clusters (**Figure 1C, S1A**). The smaller cluster is made up of the 4 most abundant lipids, Ergosterol, PC 16:1_16:1, PC 16:1_18:1, and PE 16:1_18:1 (**Figure 1C, S1A**), that together comprise around 25% of the total lipidome. The 4 lipids present in the small cluster are sufficient to identify the 4 clusters of the yeast strains, providing initial lipid signatures for each cluster (**Figure S1E**). Although this approach would need to be verified with a larger number of strains, this observation implies that the abundance of only the few most abundant lipids could already serve as biomarkers for strain identity.

### Effects of strain background on lipid class composition

To further investigate lipidome variations using an untargeted comparison, we determined which lipid species show the largest variation between the individual strains. We therefore investigated the z-scores of the individual lipid species (**Table S3**), which represents the deviation of the abundance from the overall mean abundance of the lipid species. Thus the z-score provides a metric for relative enrichment or depletion between strains. Graphical representation of the lipid z-scores per strain in a bubble plot reveals lipid features that could further distinguish the individual strains (**Supp Figure 2A**). Most notably, TAG species stood out as highly increased in RM and CEN strains compared to the other strains (**Figure 1D**), which was not readily observed from the heatmaps. Consistent with the high z-scores for nearly all TAG species in RM11 and CEN, the lipid class composition shows that TAG is the most abundant lipid in CEN.PK and RM11, at 17 mol% and 19 mol% of total lipids, respectively (**Figure 1E**, **Table 1**). This is surprising, as in early exponential growth cells are expected to mainly synthesize membrane lipids and the synthesis of storage lipids such as TAG mainly occurs in the stationary phase [14,15]. In RM11, the increased abundance of TAG occurs seemingly at the expense of mainly PI (**Figure 1D)**, which would be consistent with the balance of PI synthesis and TAG synthesis regulated by via Opi1/Ino2/Ino4 [50], also known as the Henry regulatory circuit. In CEN.PK, the elevated TAG abundance is at the expense of all three major glycerophospholipids PC, PE, and PI (**Figure 1D, S2A**).

**Figure 2.**
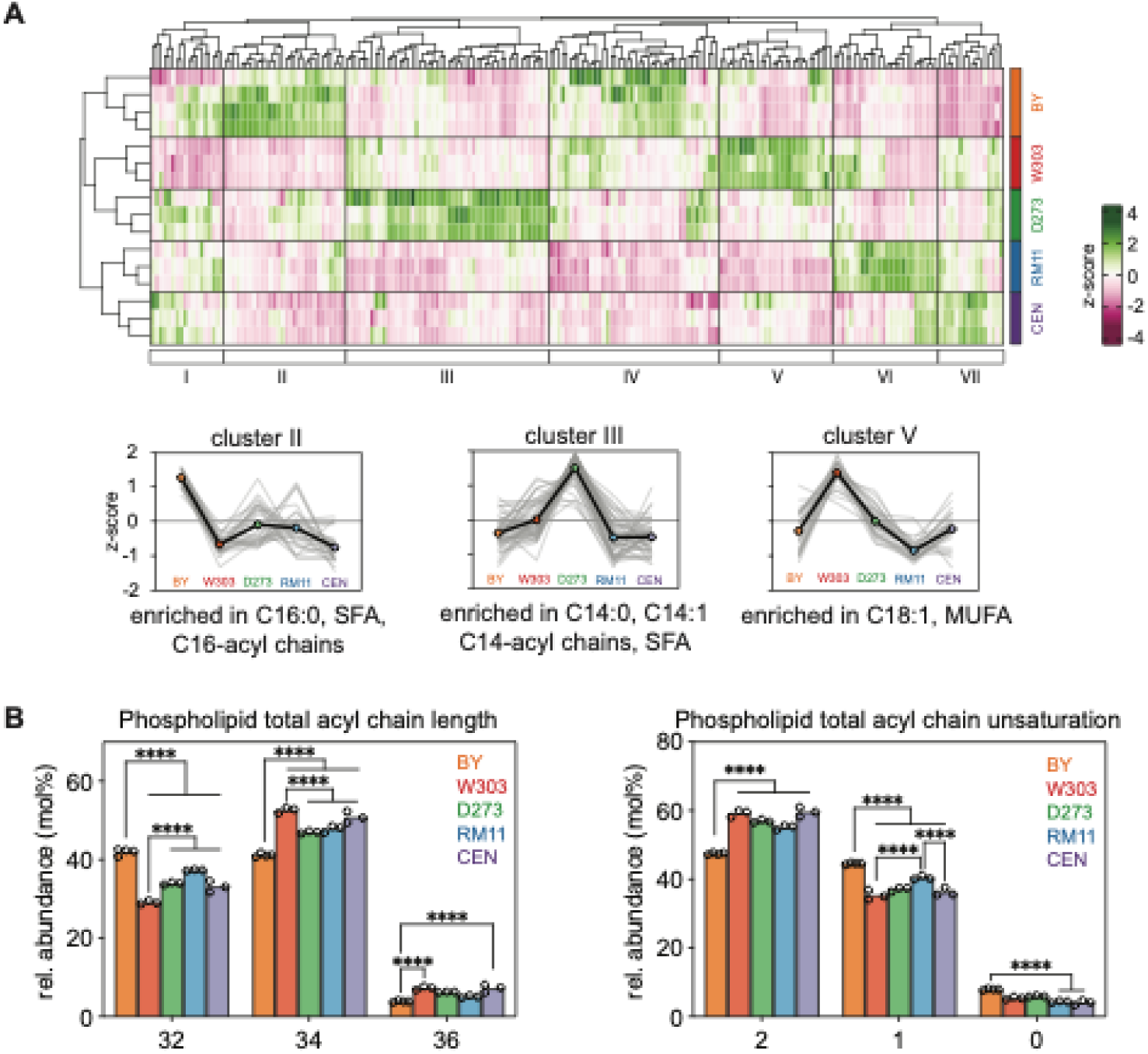
Yeast wild type strains show specific acyl chain compositions. (A) Heatmap and clustering of lipid z-scores. For selected clusters, a line-graph is added to indicate the z-scores of the lipids present in the cluster for each strain. Thin, grey lines indicate individual species, and the average z-score of the cluster is given as coloured symbols connected by a black line. (B) Total acyl chain length (left) and total acyl chain unsaturation (right) of the glycerophospholipids (PC, PE, PS, PI, PG, PI) included in the dataset. Data is depicted as the mean with individual replicates shown (n=4 for BY, n=3 for others). Stars indicate statistical significance (****: p < 0.0001) as determined by 2-way ANOVA with multiple comparisons between strains using Tukey’s method to correct for multiple testing. For clarity, selected results of statistical tests are shown..

Surprisingly, the bubble plot also revealed large variations in the z-scores for the species of the major glycerophospholipid classes PC, PE, and PI (**Figure 1D**). In example, BY shows high z-scores for most PI species and low z-scores for nearly all PC species, indicating an enrichment in PI and depletion of PC compared to the other strains. This indicates that although BY has been identified as the major glycerophospholipid class is PI [13,14], this may be strain specific. Indeed, the lipid class composition shows that while PI is the most abundant glycerophospholipid class in BY (**Figure 1E**, **Table 1**), it is PC in W303, and PE in D273. The increased abundance of PC in W303 is mainly at the expense of the anionic glycerophospholipids PI and phosphatidic acid (PA), leading to a reduction of the total amount of anionic lipids. In D273, the increased abundance of PE is at the expense of both PI and PC could create curvature stress, because PE is a cone shaped lipid that affects membrane intrinsic curvature, the strength of which depends on the acyl chains [51]. In RM11 and CEN, PC and PE are equally abundant and have the lowest abundance of PI compared to the other strains (**Figure 1E**, **Table 1)**, consistent with the negative z-scores observed for all PI species (**Figure 1D**). These observations provide handles to solve the long standing debate as to whether PI or PC is the most abundant glycerophospholipid class in yeast [13,52–54].

The z-score bubble plot revealed different trends between differences in the abundance of specific lipid classes, but for some lipid classes the species show divergent trends (**Figure 1C, S2A**). For example, in RM11 and CEN all PI lipids show negative z-scores irrespective of the lipid species, whereas in BY, W303 and D273 some species have a positive z-score where others have a negative. This is particularly the case for the species of PC and PE, which show broad distributions of z-scores in all 5 strains (**Figure 1D**). This observation indicates that while some lipid classes behave as a cohort in specific strains, for example TAG in RM and CEN, for most classes there is large variation in the behaviour of the individual species between strains.

### Effects of strain background on lipid species composition

To gain more insight into the changes in the lipid species we used clustering analysis of the lipid z-scores, which revealed 7 major clusters in the lipid z-scores heatmap that can distinguish individual strains and could be used as the lipid signatures for each strain (**Figure 2A, S3D**). We used lipid ontology (LION, [55]) enrichment analysis to identify enrichments of specific glycerolipid species or properties in each cluster. This approach not only takes into account the lipid class, but also the molecular species and for the glycerophospholipids the identity of both fatty acyl chains. We found specific acyl chain enrichments in clusters containing lipids with high z-scores for BY, W303, and D273 (Cluster II, V, III, respectively; **Figure 2A**). Cluster II, which contains lipid species with high z-scores for BY, was found to be enriched in C16:0 acyl chains specifically for saturated fatty acids (SFA) and C16 acyl chains in general (**Figure 2A**). Consistently, the glycerophospholipid acyl chain composition of BY shows strikingly higher abundance of C16:0 acyl chains compared to other strains (**Figure S2B**). The high abundance of C16:0 lipid acyl chains consequently leads to shorter and more saturated glycerophospholipid molecular species (**Figure 2B**) and similar effects were observed for TAG species (**Figure S2C**). In contrast, lipids in Cluster V are to be enriched in C18:1 acyl chains specifically and monounsaturated fatty acids (MUFA) in general (**Figure 2A**). Lipids in Cluster V show high z-scores in W303 and consistently W303 shows far higher abundance of C18:1 compared to BY (32 mol% compared to 22.5 mol%, respectively). However, this effect is only minor compared to D273, RM and particularly CEN (**Figure S2B**). Although W303 shows high z-scores for PC 18:1_18:1, PC 18:1_18:1, PI 18:1_18:1 and even LPC 18:1 and LPE 18:1, increased abundance of these lipids has only minor effect on the PL species composition (**Figure 2B**) likely due to low abundance of these species. Finally, Cluster III that contains lipids with high z-scores in D273 was found to be enriched in C14:0 and C14:1 acyl chains specifically (**Figure 2A**). Indeed, D273 shows higher abundance of C14 acyl chains in the glycerophospholipids (5.2 mol%) compared to the other strains (3.3 - 4.1 mol%, **Figure S2B**), although they are still a minor portion of the total acyl chains. The high levels of C14 acyl chains leads to an increase in C30 glycerophospholipid species but not the shorter C26 and C28 species. Interestingly, although shorter lipid species are typically mainly found in PI [51], the C14 acyl chains enriched in Cluster 3 are present in PE and PC as well as in PI (**Table S3**).

### Yeast strains show high heterogeneity in transcript abundance

The observed differences in the lipidome (**Figure 1,2**) indicate variation in the regulation of lipid metabolism between the strains. We thus set out to identify potential mechanisms that could underlie these variations. The selected yeast strains are auxotrophic for various amino acids due to the presence of auxotrophy selection markers. As the presence of auxotrophy markers has previously been shown to affect genetic interactions and cellular amino acid metabolism [56], we reasoned that they could indirectly affect lipid metabolism. This is particularly the case for *MET15*, as the Met15 protein is involved in the biosynthesis of cysteine and methionine, the latter being an upstream substrate in the synthesis of ergosterol and the *de novo* synthesis of PC. Manual comparison of the strain specific lipidomes and the auxotrophy marked present did not reveal a pattern, therefore it is unlikely that under these conditions the presence of auxotrophy markers influences the lipidome. For *MET15* specifically, this was further supported by a previous work reporting only little lipidome variation between BY4741 and BY4742, despite these strains differing in the presence of the *MET15* and *LYS2* selection markers [14]. As BY4741 and BY4742 also differ in mating type, we conclude it is also unlikely for the mating type to have an effect on the lipidome.

Although the regulation of many lipid metabolic pathways remains unknown, several mechanisms of transcriptional regulation of lipid biosynthesis have been characterized in detail [50,57,58]. We therefore used RNA sequencing to determine the transcriptome, aiming to identify differences in gene expression levels that could explain the observed differences in the lipidome. A total of 6445 mRNA transcripts could be mapped to the S288C reference genome (**Table S4**), which is over 95% of the total ORFs listed in the yeast genome database [46]. The presence and absence of strain specific markers verified the identity of the used strains (**Figure S3A**). Importantly, variation of read counts distributed heterogeneously throughout the genome, showing that there are no chromosome regions duplicated between the strains and the strains do not vary in karyotype (**Supp S3B**).

We compared the transcriptomes of the individual strains pairwise, and identified a striking total 2140 differentially expressed genes (DEGs; cutoff set at as |Log_2_FC| > 1 and p_adj < 0.05) throughout the dataset (**Figure 3, S3A,B, Table S5**) corresponding to roughly a third of the mapped transcripts. Between the individual strains, the number of DEGs ranged from 93 (D273 *vs.* W303) to 1124 (D273 *vs.* RM11). Clustering of the strains based on the transcriptome DEGs (**Figure 3**), as well as PCA and distance analysis on the total dataset (**Supp S4C,D**), showed markedly different strain clusters compared to the clustering based on the lipidome (**Figure 1C, S1C,D**). In the DEG clustering, the closest related strains are W303 and D273 and consistently comparison of these strains show the lowest number of DEGs (**Figure 3, S4B**). RM11 is most different from the other strains in terms of the transcriptome, showing the largest number of DEGs in the strain comparisons. It is most distant from D273 and W303, with 1124 and 649 DEGs, respectively, and seems to be closest related to BY while still showing 470 DEGs compared to BY. Notably, although in the lipidome analysis BY is most distant from other strains, in the transcriptome analysis BY is oriented more toward the center of the other strains (**Figure 3A, S4C,D**).

**Figure 3.**
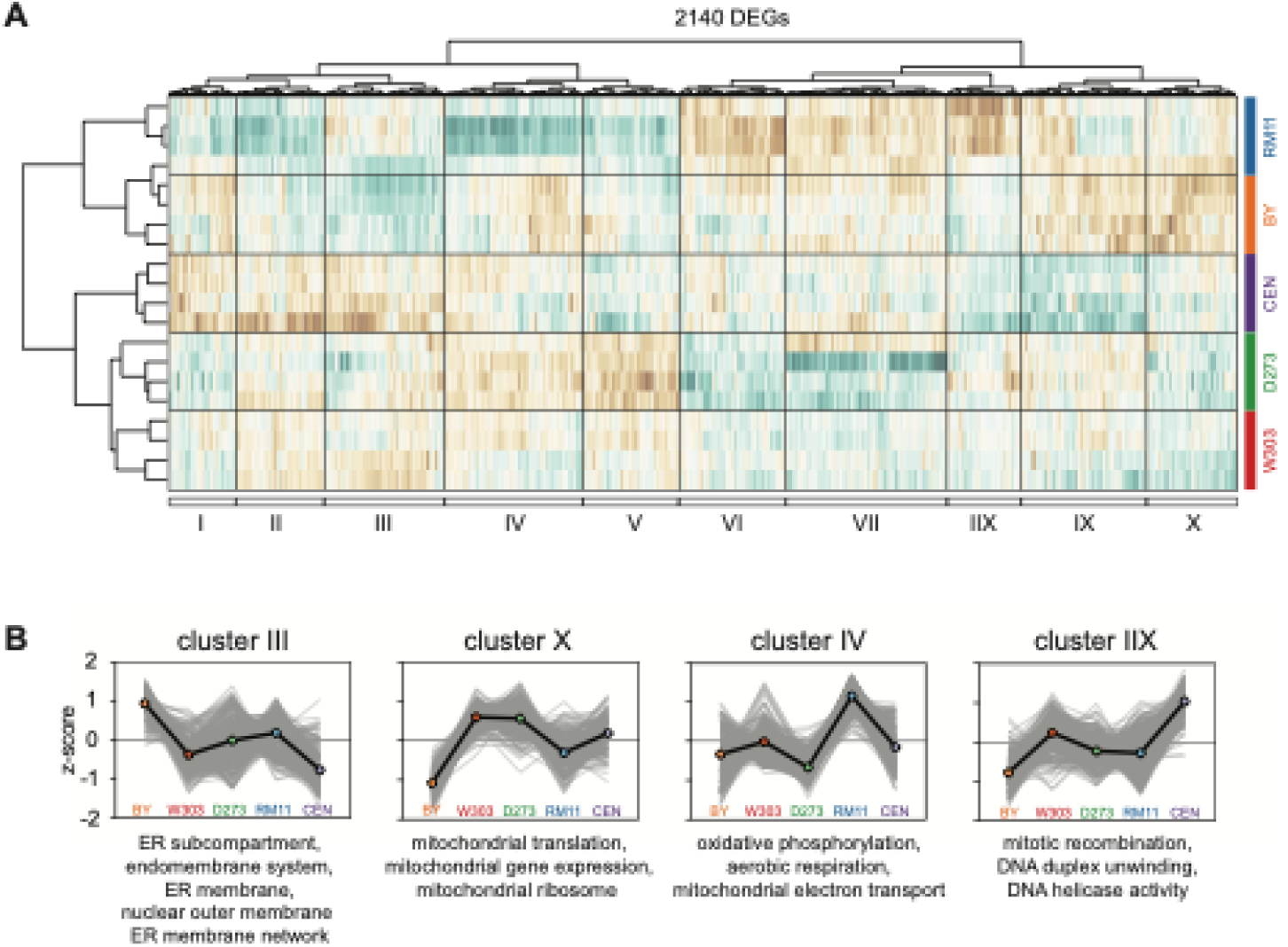
Transcriptome variation between wild type yeast strains. (A) Heatmap and clustering of the z-scores of all DEGs (2140 in total) identified in the transcriptome dataset. Underlying data can be found in Table S5. (B) For selected clusters, a line-graph is added to indicate the z-scores of the lipids present in the cluster for each strain. Thin, grey lines indicate individual species, and the average z-score of the cluster is given as coloured symbols connected by a black line. Underlying data for z-scores can be found in Table S5 and for GO-term enrichments in Table S6.

Clustering analysis of the transcript z-scores identified 10 clusters of highly correlated DEGs (**Figure 3**, **S4E**). To uncover strain-specific transcriptional patterns, we performed GO-term enrichment analysis on the genes in each cluster (**Table S6**). Notably, the BY strain showed high z-scores in Cluster III, which is enriched in GO terms related to the ER and nuclear envelope (**Figure 3**). This indicates increased transcription of genes encoding ER proteins in BY. In contrast, both W303 and D273 exhibited high z-scores in Cluster X, enriched in GO terms related to mitochondrial gene expression and protein translocation, consistent with these strains being widely used in mitochondrial research. Interestingly, BY displayed negative z-scores for DEGs in this cluster, reflecting reduced mitochondrial function and poor respiration capacity. However, despite the large variations observed in lipid composition across strains (**Figure 1, 2**), no GO terms related to lipid metabolism were enriched in any of the clusters. This suggests that the transcriptional variation underlying lipid metabolism is either diffused across many genes or regulated beyond the transcriptional level.

### Comparative transcriptome analysis indicate reduced ARE2 and HMG1 expression to underlie low EE levels in BY background

One of the most striking lipidome variations was the strong reduction in EE lipids in BY compared to the 4 other strains (**Figure 1C, 1E, Table 1**). We therefore selected this lipid phenotype as a model for the comparative analysis (**Figure 4A,B**). The reduction in EE levels is exclusive to BY, so we reasoned that transcriptome variations that underlie this phenotype must be consistent in the comparisons of BY to the other strains. We therefore determined the genes that were identified as a DEG in all 4 comparisons of BY to the other strains and that show consistent increased or decreased levels in all comparisons. We identified 31 DEGs that were consistently increased in the other strains compared to BY, and 8 genes that were decreased in the other strains compared to BY. Of these 39 identified DEGs, 1 gene was directly related to EE biosynthesis (*ARE2*) and 2 were related to ergosterol biosynthesis (*HMG1, CYB5*) (**Figure 4B,C**).

**Figure 4.**
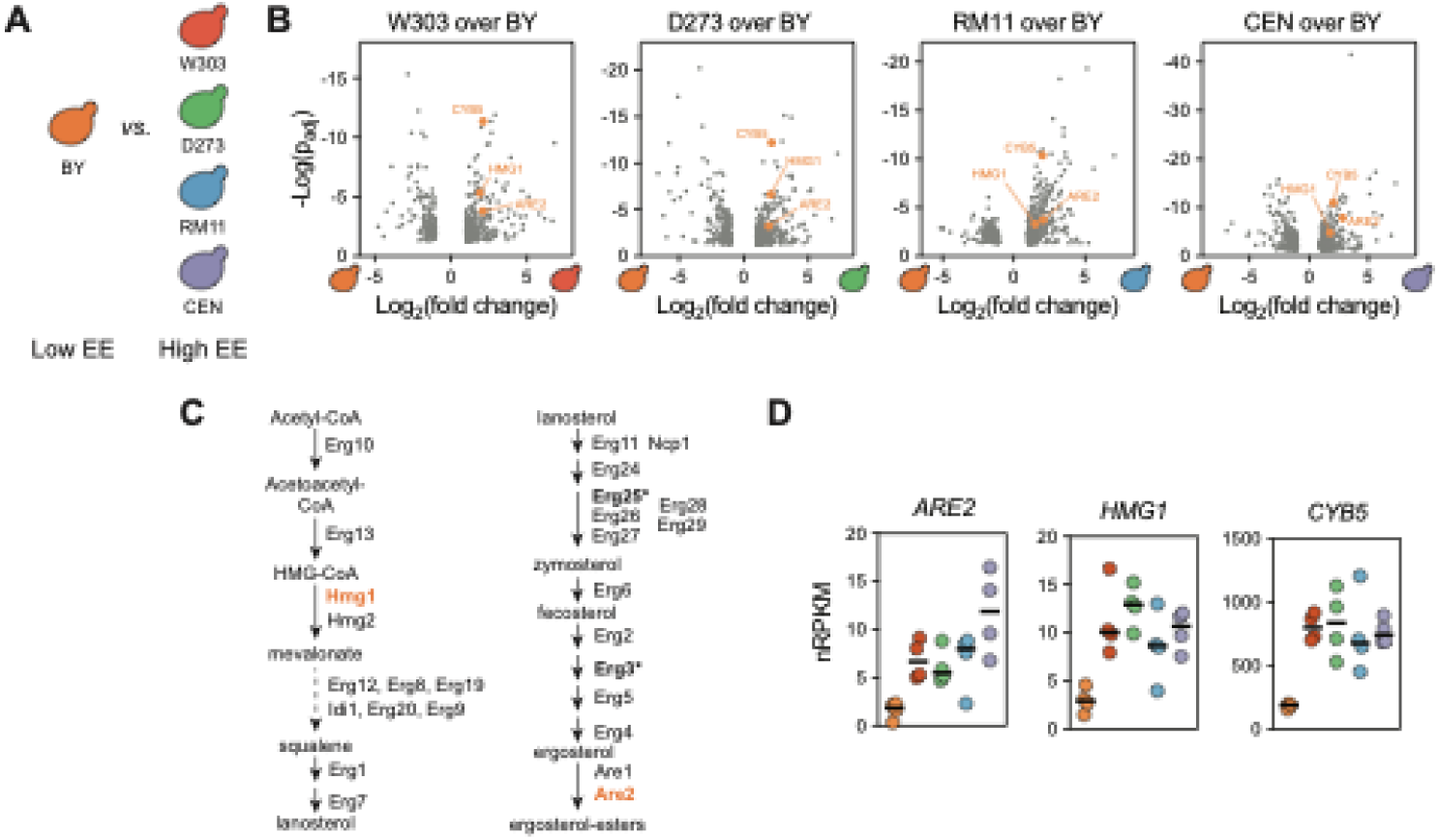
Comparative transcriptomics analysis reveals correlation of low *ARE2, HMG1,* and *CYB5* transcript levels with low EE in BY background. (A) Cartoon showing the pairwise transcriptomics comparisons of BY versus the other 4 strains. (B) Volcano plots of transcriptomics comparisons (as in Figure 3A). For clarity, only the significant hits are shown (in grey). Genes related to ergosterol esters (*ARE2, HMG2, CYB5, PLN1;* described in text) are indicated in orange. (C) Simplified overview of the lipid biosynthetic pathways involved in ergosterol ester synthesis. Proteins encoded by the identified hits *HMG1* and *ARE2* are indicated in orange. Proteins that use Cyb5 as a cofactor are indicated with an asterisk. (D) Transcript numbers (normalized reads per kilobase million) for the identified DEGs related to ergosterol ester biosynthesis.

*ARE2* encodes one of two Acyl-CoA:sterol acyltransferase enzymes (Are1 and Are2), of which *ARE2* is the major contributor to EE synthesis [59,60]. Total loss of *ARE2* reduced EE levels by over 50% [60], in agreement with the correlation between low EE levels and low *ARE2* transcript abundance in BY. However, as we find a greater than 50% reduction of EE abundance in BY compared to others, this could indicate that there are additional factors playing a role in the low EE levels in BY. We additionally identified two genes related to ergosterol biosynthesis: *HMG1*, encoding one of the enzymes involved in ergosterol biosynthesis, and *CYB*5, encoding the cytochrome b5 that acts as a cofactor to several ergosterol biosynthetic enzymes. *CYB5* encodes the yeast cytochrome b5, which acts as a heme-dependent electron donor and is a cofactor to the C-5 sterol desaturase Erg3 and the C-4 methyl sterol oxidase Erg25. A reduction in *CYB5* transcript levels could lead to a reduction of the Cyb5 cofactor, which in turn could lower the metabolic activity of Erg3 and/or Erg25. However, as neither Erg3 nor Erg25 are rate limiting enzymes, it is unlikely that a reduction in *CYB5* transcript levels would affect Erg synthesis and consequently EE levels. *HMG1* encodes one of the two HMG-CoA reductase enzymes (Hmg1, Hmg2) that catalyze the rate limiting step in ergosterol synthesis. As overexpression of *HMG1* was previously shown to increase EE levels roughly 5-fold [61], increased *HMG1* transcript levels is likely to contribute to increased EE in W303, D273, RM11 and CEN compared to BY.

## Discussion

The budding yeast *Saccharomyces cerevisiae* is one of the most widely used model organisms in biological research and has provided invaluable insights into genetics, cell biology, and metabolism [5]. Despite its status as a model eukaryote, there is no single laboratory strain or “wild type” yeast. Instead, a plethora of different laboratory strains are used across various fields and in research groups. These strains have distinct genetic backgrounds, which can lead to substantial phenotypic variation. Previous studies have described genetic background based effects mainly in the context of phenotypic traits, such as growth, stress resistance and metabolic responses [22,24,26–29,62,63]. Additionally, strain-dependent differences have been observed at the transcriptomic [26,64,65] and proteomic levels [65–68], further emphasizing the impact of genetic background on cellular processes. Although previous studies have suggested potential differences in lipid composition and lipid metabolism related phenotype between strains, a systematic analysis of how genetic background influences the yeast lipidome has been lacking. In this study, we systematically compared the lipidome of five widely used wild-type yeast laboratory strains cultured under standardized conditions, revealing striking differences in lipid class composition, total acyl chain composition, and individual lipid species. Given the extent of these lipidomic variations, we sought to identify potential regulatory mechanisms by performing transcriptomic analysis to examine strain-dependent differences in gene expression. While transcriptional variation was extensive, with a total 2140 differentially expressed genes (DEGs) identified in the dataset, we found that most of the observed differences in the lipidome could not be explained at the transcriptomic level. This suggests that regulatory mechanisms beyond gene transcription, such as regulation via protein stability, protein localization and/or enzyme activity, play a key role in shaping the yeast lipidome.

### Variation in the lipid class composition between strains

Glycerophospholipids make up the bulk of the yeast lipidome, comprising over 75% of the total membrane lipids. Whether PC or PI is the major glycerophospholipid in yeast has been under debate, and studies have reported conflicting results. The abundance of different glycerophospholipids depends heavily on the culture conditions [14,15] and may be affected by the analytical methods used [52]. We find that the identity of the major membrane lipid (PI, PC, or PE) differs between strains (**Figure 1E**, **Table 1**), which should be taken into account when selecting a strain for specific studies. In BY4741 PI was identified as the most abundant glycerophospholipid class in early exponential growth, in agreement with previous studies [14,15], whereas in W303 PC was identified as the most abundant glycerophospholipid (**Figure 1E**). It must be noted that at later points in the growth curve the level of PI in BY4741 significantly decreases, leading to PC being the major PL also in this background [14,15]. Similarly, in the presence of exogenous choline in the culture media, which allows for nett PC synthesis via the Kennedy pathway, PC can become the predominant lipid in BY [17,69,70]. In D273, we find PE to be the major glycerophospholipid (**Figure 1E**, **Table 1**), which is unexpected as PE is a type II lipid with non-bilayer propensity [51]. To maintain membrane integrity, the ratio of the non-bilayer preferring PE and the cylindrical bilayer lipid PC was proposed to be tightly maintained [71]. Increases in PE-to-PC ratio and the accumulation of PE-methylation intermediates have been implicated to cause lipid bilayer stress and activate the unfolded protein response [54,72,73]. As we do not observe transcriptome variations consistent with UPR activation in D273 it is unlikely that this increase in PE is found throughout all organelles, particularly in the ER where membrane physical properties are sensed and homeostatically regulated [74,75]. As D273 was selected for for high respiratory capacity and being abundant in mitochondria and mitochondria are highly enriched in PE [76], we speculate that mitochondrial PE is likely the origin of the increase in PE in the total lipidome. Consistently, D273 shows high expression of mitochondria related genes (**Figure 3**) compared to the other strains.

RM11 and CEN strains were found to have very similar lipidomes compared to the others and cluster closely together (**Figure 1C**). They contain high amounts of the storage lipid TAG (**Figure 1E**, **Table 1**), which in BY4741 is low abundant in early exponential growth conditions [14,15].

Though ergosterol esters were thought to be only a minor portion of the storage lipids in yeast, we found that this may be exclusive to BY as the four other strains show strikingly higher EE levels (**Figure 1E**, **Table 1**). Transcriptomic analyses identified two key lipid metabolic enzymes, *ARE2* and *HMG2*, with lower expression in BY. These observations argue that BY is not a suitable strain to study EE metabolism, in contrast to W303 and D273 in which TAGs and EEs are equally abundant.

### Yeast strains differ in lipid acyl chain length and unsaturation

When investigating the molecular species composition of membrane lipids, we observed large variation in total acyl chain length and unsaturation across strains (**Figure 2B, S2B,C**). Analysis of the glycerophospholipid acyl chain composition confirmed that in BY, over 65% of the acyl chains are composed of C16:0 and C16:1 (**Figure S2B**), consistent with previous reports on the total acyl chain composition [49,51,70]. Compared to the other four strains, BY exhibited a clear shift toward shorter and more saturated acyl chains, predominantly at the expense of C18:1.

Membrane lipid unsaturation in yeast is controlled by the Δ9 acyl-CoA desaturase *OLE1*, which is transcriptionally regulated by the Mga2/Spt23 system in response to membrane properties [77,78]. Since *OLE1* was not identified as a DEG in any of the comparisons, transcriptional regulation of fatty acid desaturation does not explain the observed variation in acyl chain composition. Moreover, when comparing BY to the other four strains, none of the consistently upregulated or downregulated DEGs were directly related to fatty acid synthesis, elongation, or desaturation. Thus, the differences in acyl chain composition across strains must arise from regulatory mechanisms beyond transcriptional control, such as post-translational modifications, enzyme activity regulation, or differences in the metabolic flux.

### Transcriptomic variation only explains lipidome heterogeneity to small extent

Transcriptome profiling identified large variation in transcript abundance was observed between the 5 different wild type strains (**Figure 3, S4A,B**), with a total of 2140 DEGs identified (**Figure 3**). Although 113 DEGs were annotated with the GO-term ‘lipid metabolic process’, this go term was not significantly enriched in the total dataset or any of the identified transcript clusters and manual analysis of the lipid metabolic pathways neither reveal the up- or downregulation of genes related to specific lipid metabolic pathways. This indicates that the majority of the observed lipidome heterogeneity is not created by changes in regulation at the level of gene expression and for most observed changes in the lipidome the direct cause remains unresolved. There are various means of regulation of metabolism beyond transcriptional regulation that would not be observed in transcriptomics analysis, such as post-translational modifications, protein stability and degradation, protein localisation, and substrate availability. Future studies integrating proteomics, phosphoproteomics, and organelle-specific lipidomics will be crucial to uncovering the mechanisms underlying the regulation of lipid metabolism.

Another important limitation of the transcriptomics analysis is the large number of genes of unknown function in *S. cerevisiae.* Of the 6776 open reading frames (ORFs) reported in the SGD, 941 (13.9%) are annotated as ‘proteins of unknown function’. Among the 2140 DEGs identified in our dataset, 327 genes (15.3%) fall into this category, and 388 DEGs (18.1%) lack a standardized gene name. Given this substantial proportion of poorly described genes, it is likely that some of these genes are involved in lipid metabolism or its regulation, but their roles remain uncharacterized. Additionally, because our mRNA sequencing data were aligned to the S288C reference genome, strain-specific ORFs that are absent from S288C but present in other yeast genomes were not captured. These missing annotations may contribute to the unexplained lipidomic variation observed between strains. Future studies using functional genomics approaches to compare different genetic backgrounds to the BY-background may help uncover novel regulators of lipid metabolism encoded within these poorly characterized regions of the genome.

### Concluding remarks and perspectives

Our findings highlight that genetic background plays a major role in shaping the yeast lipidome, emphasizing the importance of strain selection in lipid metabolism studies. The substantial lipidomic variation observed between widely used laboratory strains suggests that strain-specific variations in the regulation of lipid metabolism could underlie the observed variations in lipid metabolism related phenotypes [31,32]. As a result, studies performed in a single strain may not always be directly translatable to others, and strain background should be carefully considered when interpreting lipid-related phenotypes. In example, it remains to be seen if the patterns of lipidome adaptation in response to culture conditions and growth phase, which are well established in BY [14,15], occur in similar fashion in other strains. This would be particularly interesting for non-laboratory strains such as wine and beer yeasts, natural isolates and clinical isolates, that show genetic variation from the reference far beyond that of laboratory strains [21,22]. Beyond bulk lipidomic differences, subcellular lipidomes may also vary between strains, influencing organelle function, membrane properties, and cellular adaptation to environmental conditions. For organelle-specific lipid analysis typically strains are selected in which the organelle of interest is highly abundant [17,79–82], which may have an underestimated role in determining the outcome of the analysis due to strain-dependent lipidome variations.

Comparative studies using multiple yeast strains provide a unique opportunity to identify novel regulators of metabolism. By analyzing strains with distinct lipidomic profiles or lipid metabolism related phenotypes it becomes possible to uncover genetic factors (quantitative trait loci) that regulate lipid metabolism. For example, segregant analysis from a cross of BY4741 and the wine yeast AWRI1631 revealed a relation of *PIG1, PHO23,* and *RML2* to neutral lipid abundance [33]. Recent advances in genetic technologies such as CRISPR/Cas9 and transposon based screening [83,84] expand genome wide studies to other strains beyond BY, providing opportunities for comparative genetic screening approaches at large scale. Such approaches broaden the perspectives of the use of yeast as a model system and may in the future help to bring the yeast genome to a full annotation of genes and ORFs.

## Methods

### Yeast strains and culture conditions

BY4741 (EuroSCARF Y00000; *MATa his3Δ1 leu2Δ0 met15Δ0 ura3Δ0*) and CEN.PK2-1C (EuroSCARF Y30000A; *MATa his3Δ1 leu2-3_112 ura3-52 trp1-289 MAL2-8c SUC2*) were newly purchased prior to this study (June 2022) from EuroSCARF (Oberursel, Germany). RM11-1A (*MATa leu2Δ0 ura3Δ0 HO::kanMX*) and D273-10B (*MATα mal*) were kindly provided by Dr. Karin Aethenstadt (Universität Graz, Graz, Austria). The *ADE2*-corrected strain variant of W303-1A (*MATa leu2-3,112 trp1-1 can1-100 ura3-1 his3-11,15*) was kindly provided by Dr. Sebastian Schuck (Heidelberg University, Heidelberg, Germany).

Strains were cultured in synthetic complete dextrose (SCD, [85]), containing per liter: 1.7 g yeast nitrogen base without amino acids and without ammonium sulphate (ForMedium), 5 g ammonium sulphate (Carl Roth), 0.79 g complete supplement mixture (ForMedium) and 20 g glucose (Carl Roth). All ingredients were added from filter sterilized stock solutions. SCD media contains 11 µM inositol [32,85] and is devoid of choline. Absence of choline in the media was verified using a choline auxotroph strain [86].

Strains were cultured to early exponential growth as described before [16,17]. Briefly, single colonies were inoculated into 3 mL SCD pre-cultures and incubated overnight (30°C, 220 rpm) until stationary. Main cultures were inoculated at OD = 0.1 in pre-warmed media and cultured (30°C, 220 rpm) to easily exponential growth (OD 0.8 - 1.1). Culture density (as turbidity) was measured as optical density at 600 nm (OD_600_) on a Ultrospec 10 cell density meter (Harvard Biochrom). When cells reached the required OD, cells were harvested by centrifugation (3300 g, 5 min, 4°C), washed with cold 155 mM ammonium bicarborbonate (AMBIC), and frozen in liquid nitrogen. Cell pellets were stored at -80°C.

For lipidomics analysis, cells were lysed by bead beating. Briefly, cell pellets corresponding to 20 OD were thawed on ice and resuspended in 1 mL 155 mM AMBIC. Cell suspension was transferred to a crew cap tube containing 0.5 mL zirconia beads and cells were subjected to bead beating 5 times for 30 seconds using a FastPrep-24 (MP Biomedicals). Cells were cooled in an ice-water bath for one minute between bead beating steps. After lysis, the lysate was transferred to new tubes (Eppendorf) and snap-frozen in liquid nitrogen. Lysates were stored at -80°C for storage or on dry ice for shipping.

### Lipid extraction and mass spectrometry lipidomics analysis

Shotgun mass spectrometry based lipidomics was performed by Lipotype GmbH as described before [13,14,16,17]. Lipids were extracted using the two-step chloroform/methanol procedure [13]. Samples were spiked with an internal standard mixture containing: CDP-DAG 17:0/18:1, ceramide 18:1;2/17:0 (Cer), diacylglycerol 17:0/17:0 (DAG), lyso-phosphatidate 17:0 (LPA), lyso-phosphatidylcholine 12:0 (LPC), lyso-phosphatidylethanolamine17:1 (LPE), lyso-phosphatidylinositol 17:1 (LPI), lyso-phosphatidylserine 17:1 (LPS), phosphatidate 17:0/14:1 (PA), phosphatidylcholine 17:0/14:1 (PC), phosphatidylethanolamine 17:0/14:1 (PE), phosphatidylglycerol 17:0/14:1 (PG), phosphatidylinositol 17:0/14:1 (PI), phosphatidylserine 17:0/14:1 (PS), ergosterol ester 13:0 (EE), triacylglycerol 17:0/17:0/17:0 (TAG), stigmastatrienol, inositolphosphorylceramide 44:0;2 (IPC), mannosyl-inositolphosphorylceramide 44:0;2 (MIPC) and mannosyl-di-(inositolphosphoryl)ceramide 44:0;2 (M(IP)2C). After extraction, organic phases were dried in a speedvac concentrator. Lipid films were resuspended in 7.5 mM ammonium acetate in chloroform/methanol/propanol (1:2:4, V:V:V) for the first phase or 33% ethanol solution of methylamine in chloroform/methanol (0.003:5:1; V:V:V) for the second phase. All liquid handling steps were performed using Hamilton Robotics STARlet robotic platform with the Anti Droplet Control feature for organic solvents pipetting.

Samples were analyzed by direct infusion on a QExactive mass spectrometer (Thermo Scientific) equipped with a TriVersa NanoMate ion source (Advion Biosciences). Samples were analyzed in both positive and negative ion modes with a resolution of R=280,000 (at m/z 200) for MS^1^ and R=17,500 (at m/z = 200) for MS^2^ experiments, in a single acquisition. MS^2^ analysis was triggered by an inclusion list encompassing corresponding MS mass ranges scanned in 1 Da increments [87]. Both MS^1^ and MS^2^ data were combined to monitor EE, DAG, and TAG ions as ammonium adducts; PC as an acetate adduct; and CL, PA, PE, PG, PI, and PS as deprotonated anions. MS^1^ only was used to monitor LPA, LPE, LPI, LPS, IPC, MIPC, M(IP)_2_C as deprotonated anions; Cer and LPC as acetate adducts and ergosterol as protonated ion of an acetylated derivative [88].

MS data was analyzed with in-house developed lipid identification software based on LipidXplorer [89,90]. Only lipid identifications with a signal-to-noise ratio >5, and a signal intensity 5x higher than in corresponding blank samples were considered for further data analysis.

### Statistical analysis of Lipidomics data and lipid ontology analysis using LION/web

All data processing was done in R (RStudio Version 2024.09.0+375). Statistical testing was done using R or Graphpad Prism (version 10.4.0).

Clusters of lipid z-scores (Figure 2) were further analysed using LION/web [55] using the targetlist mode. A targetlist was manually constructed for all glycerolipids (PL, lysoPL, DAG, TAG) present in the clustering analysis following the input requirements. To conform to the input requirements, hydroxylation annotation was removed and knowledge on the stereospecificity of the acyl chains was assumed (in example PC 16:0;0_16:1;0 was converted into PC(16:0/16:1)).

### RNA isolation, library preparation and sequencing

RNA was isolated from cell pellets using Direct-zol™ RNA Miniprep kit (Zymo research #R2051) according to manufacturer instructions. In brief, cells were lysed using Trizol® and DNA was digested using provided DNAse I. After three washing steps, the RNA was eluted. RNA concentration was measured with a NanoDrop™ (ThermoFisher) and stored at -80 °C or used immediately for library preparation.

Sequencing libraries were created using the Smart-seq2 protocol [91]. In brief, 100 ng of RNA were used for reverse transcription and subsequent preamplification PCR. PCR product was purified using Ampure XP beads at 1:0.8 ratio. Concentration of purified cDNA was measured using Qubit (Thermofisher), and quality was checked using Qsep Bio-Fragment Analyzer (NIPPON Genetics). 4 ng of cDNA was tagmented using Tn5 (Nextera) barcoded and PCR amplified. The PCR product was purified with Ampure XP beads at a ratio of 1:0.9. Library concentration was measured using Qubit and quality was checked using Qsep Bio-Fragment Analyzer. Libraries were sequenced on an Illumina Nextseq 500 at 75 bp single-end reads.

### RNA-seq data processing

Raw sequencing data in fastq format were processed with the nf-core/rnaseq pipeline v3.12.0 [92] using Star-Salmon against the S288C yeast genome assembly and gene annotations based on Ensembl v109. Expression values were then imported into R and processed with DESeq2 v1.44.0 [93]. Genes were deemed differential when they had Benjamini-Hochberg-adjusted p-value < 0.05 and an effect size of |log2FC| > 1. GO-term enrichment and pathway enrichment was done using AllianceMine via alliancegenome.org.

All RNA-seq data have been uploaded to GEO and are available under the accession number GSE289029.

## Author contributions

MFR: Conceptualization, Investigation, Resources, Formal Analysis, Data Curation, Validation, Visualization, Methodology, Software, Project administration, Supervision, Funding acquisition, Writing - original draft, Writing - review and editing; RB: Investigation, Formal Analysis; CK: Data Curation, Formal Analysis, Validation, Methodology; TH: Investigation, Formal Analysis, Data Curation, Validation, Visualization, Software, Methodology; JSH: Methodology, Validation, Resources; RE: Conceptualization, Validation, Supervision, Resources, Funding acquisition, Writing - review and editing.

## Acknowledgements

We thank Dr. Karin Aethenstadt and Dr. Sebastian Schuck for sharing yeast strains. This work was funded by the European Research Council under the European Union’s Horizon 2020 research and innovation program (grant agreement no. 866011) to R.E. and the Saarland University Start-Up funding program (“Anschubfinanzierung”) to M.F.R. RNA sequencing was performed in the Epigenomics Sequencing Facility at Saarland University.

## Supplementary Materials

**Figure S1.**
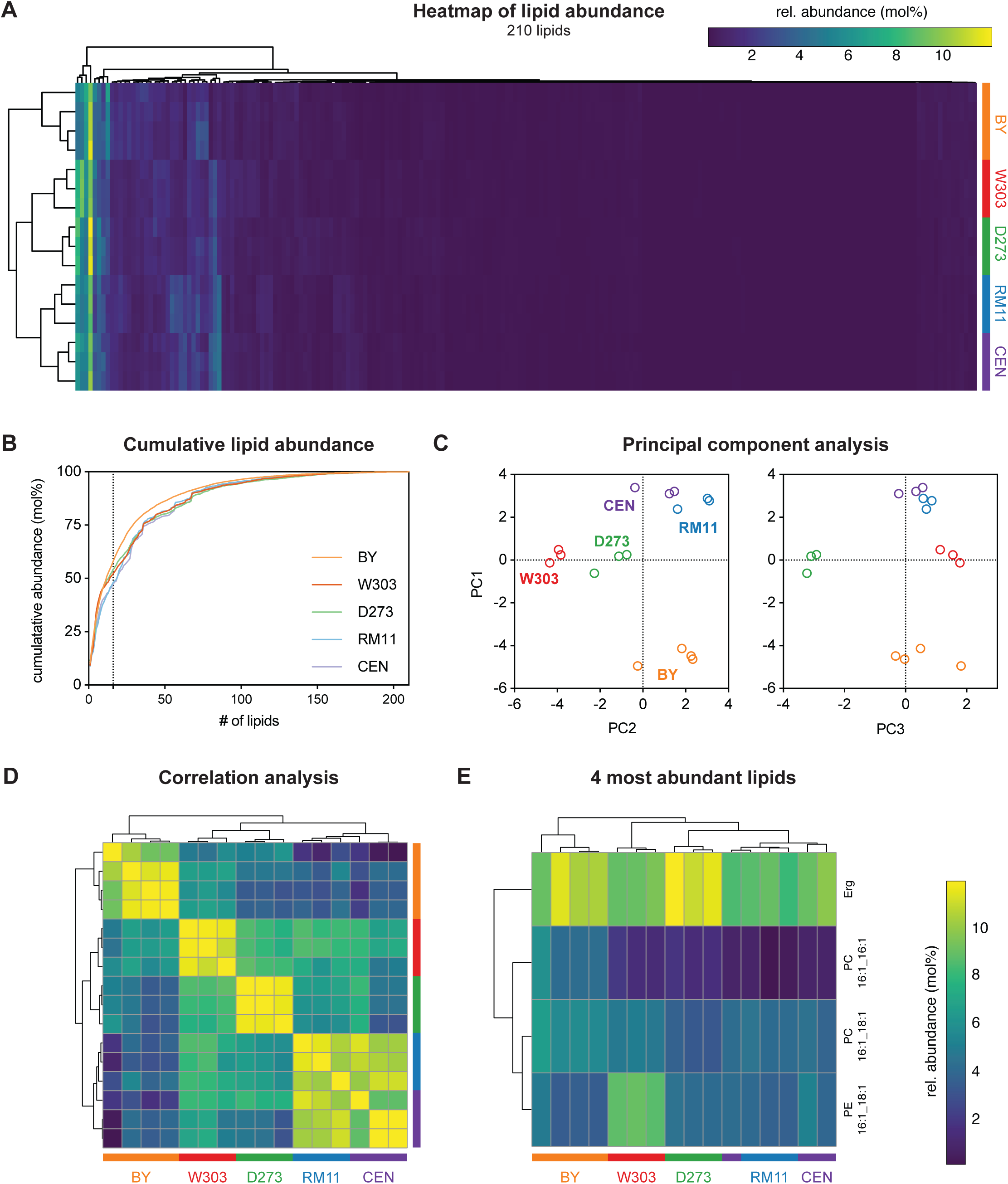
corresponding to Figure 1. A. Heatmap of the total lipid composition (210 lipid species) after lipid selection criteria. Corresponding data can be found in Table S2. B. Cumulative abundance of the lipid species in all samples. Dotted line indicates the 16 most abundant lipid species. C. Principal component analysis of the total lipidome. D. Correlation analysis (Pearson) of the total lipidome. E. Clustering analysis of the samples based on the top 4 most abundant lipid species.

**Figure S2.**
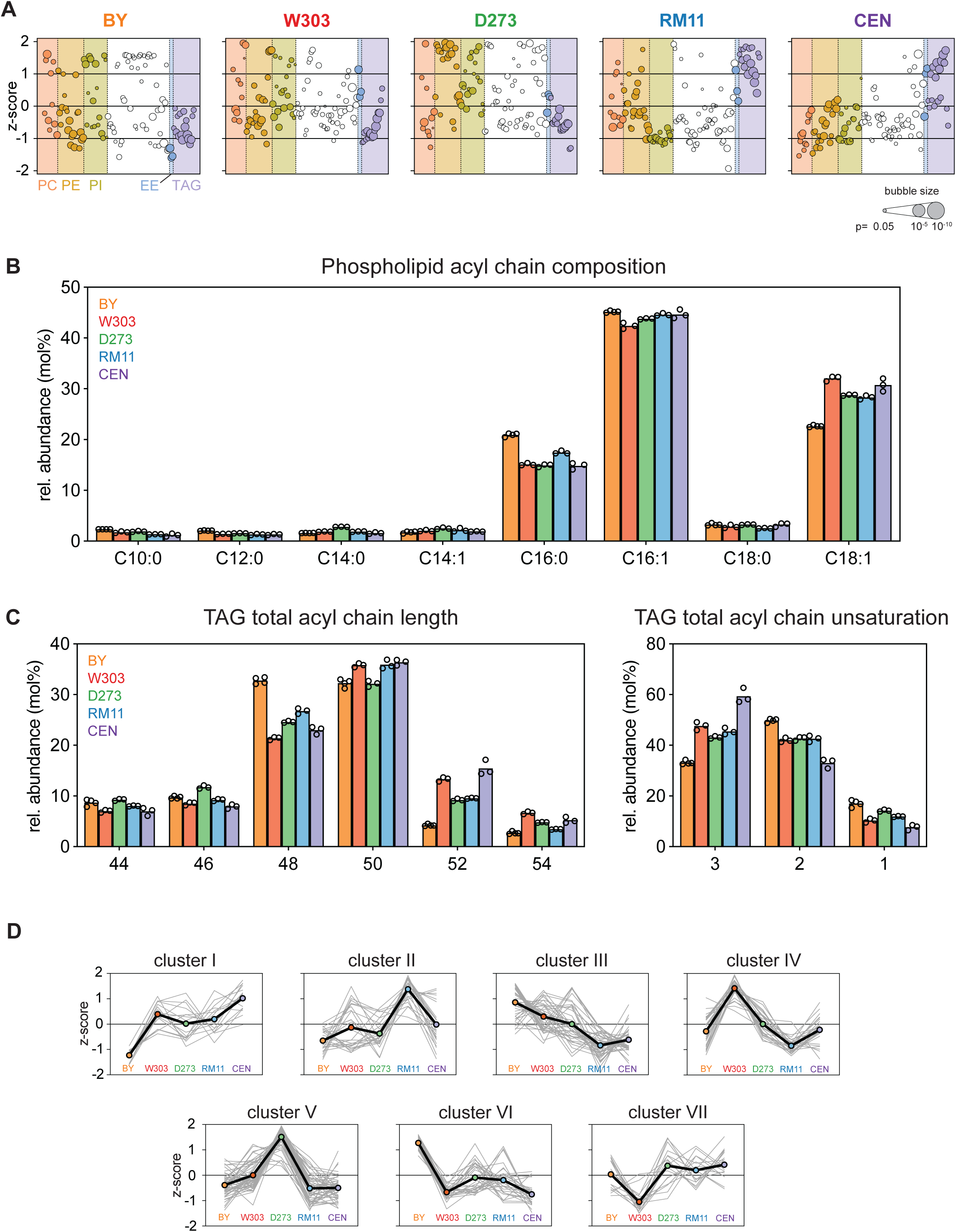
corresponding to Figure 1 and 2. A. Bubble plot of the lipid z-scores per sample. Lipid classes corresponding to Figure 1D are indicated in colored fields. Bubble size indicates adjusted p-value (AVONA with correction for multiple testing). For clarity, only species with an adjusted p-value of <0.05 are shown. B. Phospholipid total acyl chain composition. For clarity, minor acyl chain species are omitted. C. Triacylglycerol species total acyl chain length and unsaturation. For clarity, only major acyl chain lengths are shown. D. Lipid z-scores for all clusters identified in Figure 2A.

**Figure S3.**
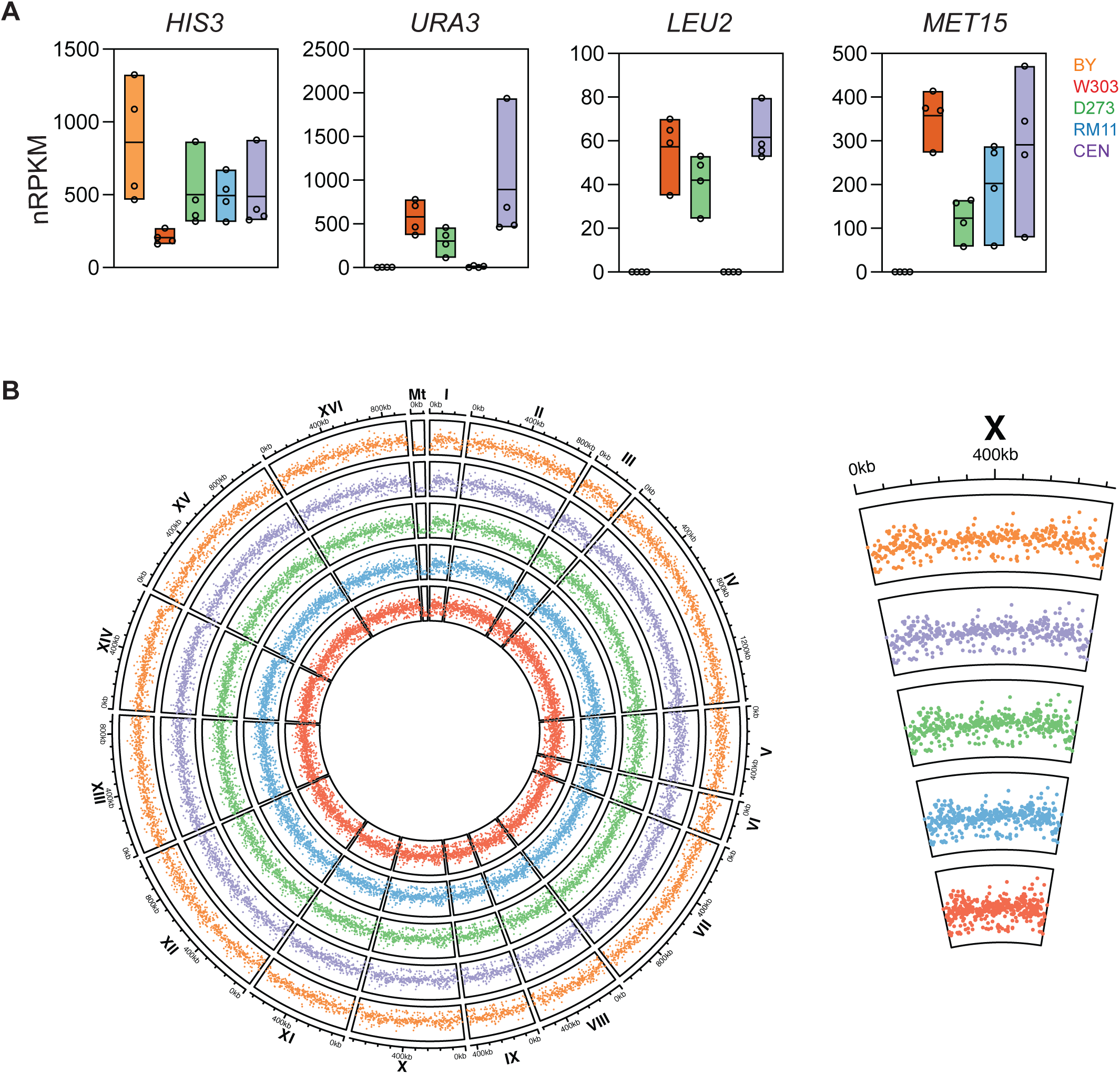
corresponding to Figure 3. A. Transcript abundance (nRPKM) for auxotrophic marker genes per strain. B. Average transcript abundance per strain for each chromosome and mitochondrial genes

**Figure S4.**
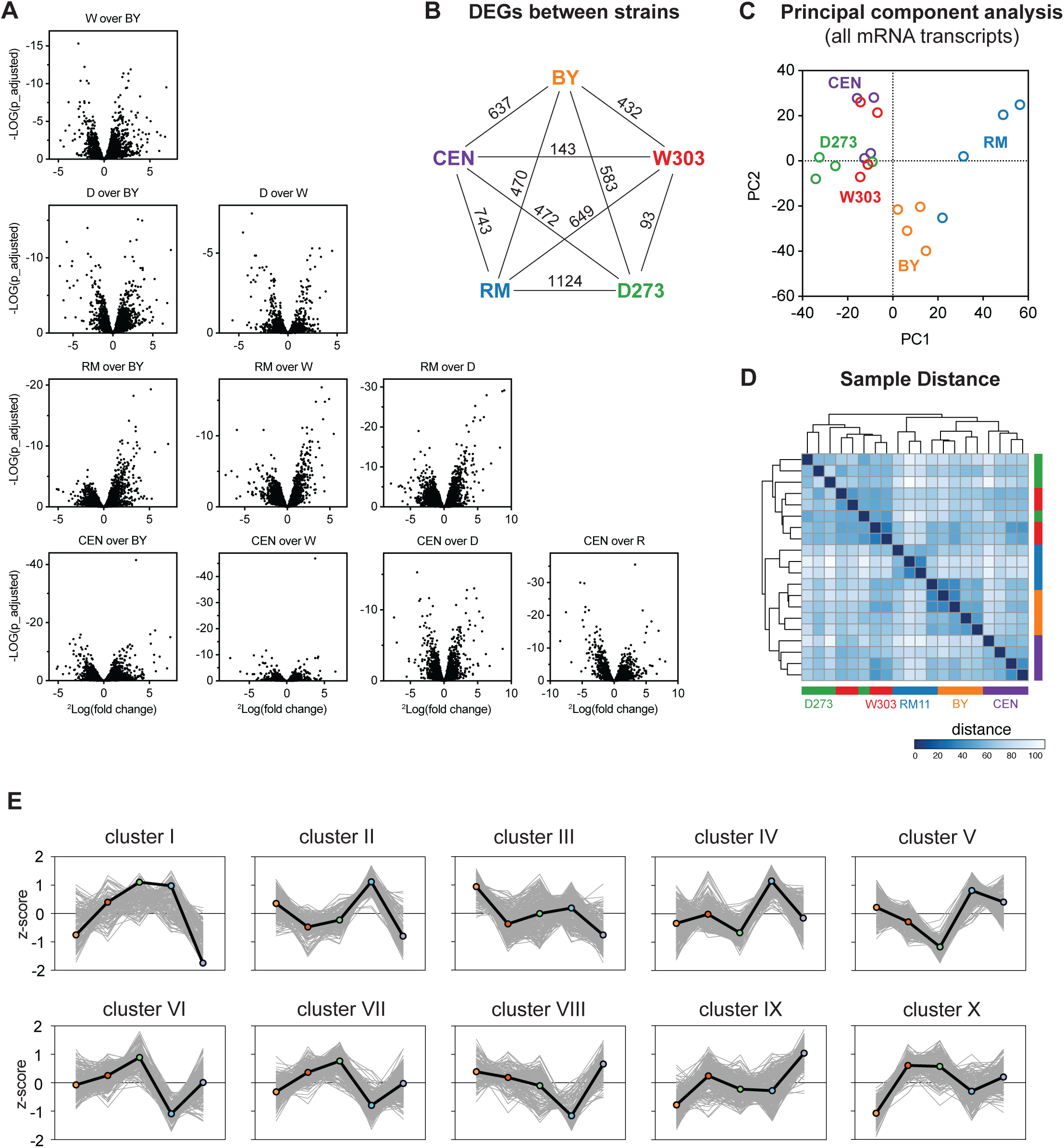
corresponding to Figure 4 and 4. A. Volcano plots of all pairwise strain transcriptome comparisons. B. Graphical overview of the number of DEGs identified in each strain comparison. C. Principal Component Analysis (PCA) based on the total transcriptomics dataset. D. Sample distance analysis based on the total transcriptomics dataset. E. DEG z-scores for all clusters identified in Figure 3A.

## Notes

https://www.ncbi.nlm.nih.gov/geo/query/acc.cgi?acc=GSE289029

### Competing Interest Statement

CK declares competing financial interests related to the publication of this
study, including being chief technology officer at Lipotype GmbH, Dresden. The
remaining authors declare no competing interests.

